# Rigid-body fitting to atomic force microscopy images for inferring probe shape and biomolecular structure

**DOI:** 10.1101/2021.02.21.432132

**Authors:** Toru Niina, Yasuhiro Matsunaga, Shoji Takada

## Abstract

Atomic force microscopy (AFM) can visualize functional biomolecules near the physiological condition, but the observed data are limited to the surface height of specimens. Since the AFM images highly depend on the probe tip shape, for successful inference of molecular structures from the measurement, the knowledge of the probe shape is required, but is often missing. Here, we developed a method of the rigid-body fitting to AFM images, which simultaneously finds the shape of the probe tip and the placement of the molecular structure via an exhaustive search. First, we examined four similarity scores via twin-experiments for four test proteins, finding that the cosine similarity score generally worked best, whereas the pixel-RMSD and the correlation coefficient were also useful. We then applied the method to two experimental high-speed-AFM images inferring the probe shape and the molecular placement. The results suggest that the appropriate similarity score can differ between target systems. For an actin filament image, the cosine similarity apparently worked best. For an image of the flagellar protein FlhA_C_, we found the correlation coefficient gave better results. This difference may partly be attributed to the flexibility in the target molecule, ignored in the rigid-body fitting. The inferred tip shape and placement results can be further refined by other methods, such as the flexible fitting molecular dynamics simulations. The developed software is publicly available.

**Author Summary:** Observation of functional dynamics of individual biomolecules is important to understand molecular mechanisms of cellular phenomena. High-speed (HS) atomic force microscopy (AFM) is a powerful tool that enables us to visualize the real-time dynamics of working biomolecules under near-physiological conditions. However, the information available by the AFM images is limited to the two-dimensional surface shape detected via the force to the probe. While the surface information is affected by the shape of the probe tip, the probe shape itself cannot be directly measured before each AFM measurement. To overcome this problem, we have developed a computational method to simultaneously infer the probe tip shape and the molecular placement from an AFM image. We show that our method successfully estimates the effective AFM tip shape and visualizes a structure with a more accurate placement. The estimation of a molecular placement with the correct probe tip shape enables us to obtain more insights into functional dynamics of the molecule from HS-AFM images.

## Introduction

Virtually all the cellular phenomena are accomplished by functional dynamics of individual biomolecules, and thus observation of their movements at the single-molecule level is a key step to understand molecular mechanisms. Among many single-molecular observation techniques(1) such as fluorescence resonance energy transfer microscopies(2–4), single particle tracking(5), optical tweezers(6), and atomic force microscopies (AFM)(7), the high-speed AFM (HS-AFM) is unique in that it can observe real-time structural dynamics of biomolecules at work near the physiological condition(8,9). The HS-AFM has been extensively used in biophysics to successfully observe structural dynamics of molecular motors(10–12), large biomolecular complexes(13,14), intrinsically disordered proteins(15), and so on. Thanks to recent developments, the frame acquisition rate of the HS-AFM reaches tens of frames per second, keeping its high spatial resolution, ~2 nm in lateral direction and ~0.15 nm in vertical direction to the stage(9).

Although the AFM is powerful, it remains difficult to estimate three-dimensional molecular structure directly from the AFM image. This is in part because the AFM is limited to the two-dimensional surface height information detected via the force to the probe. Moreover, to transform the AFM data into threedimensional structural information, one needs to know the probe tip shape, which is normally unknown *a priori*. There are several works that estimate the orientation and location of a molecule from an AFM image(16–20). In contrast, estimation of the shape of the tip used in the experiments is limited (21–24). The blind tip estimation method can infer the outer bound on the tip geometry from an AFM image without assuming any tip shape, but is somewhat sensitive to the noise in the image (25). Alternatively, Trinh and colleagues(25) developed a method to estimate the probe tip radius by using tobacco mosaic virus (TMV) as a calibrator and proposed to always observe it with the samples of interest. Although TMV is a stable and easy-to-use calibrator, still it is not always easy to observe it at the same condition as other molecules that generally requires different conditions to be stabilized on the AFM stage. Also, a recent work(26) that reconstructs the three-dimensional structures of amyloid fibril developing a method to refine the topography carefully estimates the probe tip radius. However, they use the ideal corkscrew symmetry in the specimen and it is not always applicable to other biomolecules. In general, the probe tip shape varies sample by sample. Its radius is typically sharpened to ~5 nm and sometimes less than 1 nm, and it is almost impossible to observe the shape of the probe tip at such a high resolution before using it in the experiment(8). Recently, we developed a computational method(27) that enables to reconstruct a molecular model from an AFM image taking the conformational change into account, inspired by the flexible fitting method for cryo-electron microscopy (cryo-EM) electron density map(28). We also showed that, from the result of the flexible fitting, we can estimate the effective tip radius(29). However, the biased molecular dynamics is rather time-consuming method that requires large computational resources. Thus, it is not yet useful enough to estimate the tip probe. Also, the information that can be reconstructed from the flexible fitting is limited to the parameter used in the biasing potential and sometimes it might be hard to interpret as the physical shape of the probe.

In this paper, we developed a rigid-body fitting method to simultaneously search the probe tip shape and the structure placement best-fit to a given AFM image. The tip shape can be estimated from the optimization of the similarity between an experimental AFM image and pseudo-AFM image generated from a molecular model with a varying geometry of the probe tip, as in our previous approach(29). In contrast to the previous approach, our current method directly models the physical shape of the probe tip, and thus is more straightforward to interpret the estimation results. First, by twin experiments, we examined four different image similarity scores for three monomeric proteins and investigated the relationships between the size of proteins, the image similarity scores, and the accuracy of tip shape inference. Next, we performed a twin experiment using a model of actin filament and confirmed that our method works well even when the template structure fits only to a part of an observed AFM image. Finally, we applied our method to two real experimental HS-AFM images, the actin filament and the flagellar protein FlhA_C_ ring, and estimated the probe shape in the experimental measurements.

## Methods

### Collision-detection Method for Pseudo-AFM Image Generation

We utilized the conventional collision-detection method to generate pseudo-AFM images from a structural model of the target molecule(27,30–32). The method calculates the height at which the probe hits an atom in the target molecules tethered on the stage. The stage surface is set to the *XY*-plane, on which the molecules are placed to the positive *Z* side. We approximate the probe tip as a hemisphere (the radius 0.5 to 5.0 nm) combined with a circular frustum of a cone (the half-apex angle 5 to 30 degree), with its symmetric axis parallel to *Z*-axis (Fig 1A). For each pixel, the probe center is fixed to the *XY*-coordinates of the center of the pixel. Moving down the probe from the top, the method detects the collision between the probe tip and atoms in the target. In the twin-experiments, when we make a reference AFM image that serves as an “experimental AFM image”, after determining the collision heights of all the pixels, we added spatially independent Gaussian noise with the mean 0 nm and the standard deviation 0.3 nm, which is a typical level of noise as is shown in Results.

**Fig 1.**
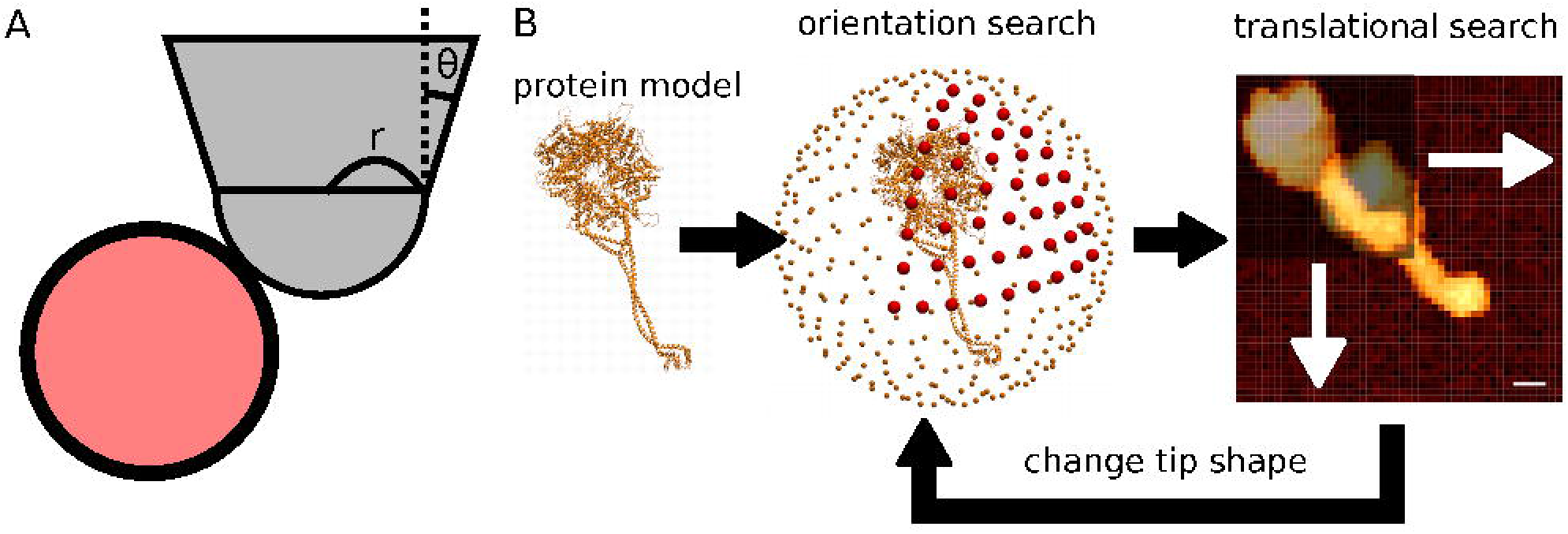
A pseudo-AFM image generation scheme and the search algorithm. (A) The AFM probe tip (grey) is modeled as a hemisphere connected to a circular frustum of a cone. The shape of tip is determined by the tip radius (*r*) and a half-apex angle of cone (*θ*). The pseudo-AFM image represents the height of the tip when the tip collides with any atom (red circle) of the target molecule. (B) The exhaustive search finds the best match in the discrete space made of the orientation and the translation of the target molecule and the radius and the half-apex angle of the probe tip.

### Image Similarity Calculation

To calculate similarity between a pseudo-AFM image generated from a molecular model and the reference image (the AFM experimental data in normal use and a generated image in the case of twin-experiments), we examined four different similarity score functions; the cosine similarity, the correlation coefficient, the pixel-RMSD, and the penalty function. It should be noted that, even though we call the AFM data “the AFM image”, it is merely the two-dimensional array of the molecular height. Therefore, the mapping to colors does not affect the analysis. Through twin-experiments, we compared their performances.

Since the experimental/reference AFM image contains non-negligible noise, the search method may tend to place the molecule at a high-noise spot as an artifact. To reduce this unfavorable case, we limited the pixels used to calculated the similarity functions. In the similarity calculation, we trimmed both pseudo- and reference AFM images by the minimum bounding rectangle of non-zero-height pixels in the pseudo-AFM image. We note that, due to the probe tip size, the pseudo-AFM image can have non-zero height even at the pixel where the molecule is absent. This is especially significant when the target molecule is tall, the probe tip radius is large, and the probe apex angle is large.

#### Cosine Similarity

The cosine similarity is the function used in our previous research of flexible fitting method(27), though it was called a “modified correlation coefficient” previously, and also used in the field of cryo-EM flexible fitting(28). The form of the function resembles to the correlation coefficient, but each term does not subtract the mean values. Thus, unlike the correlation coefficient, the cosine similarity depends not only on the relative height of pixels, but also the absolute height of pixels.

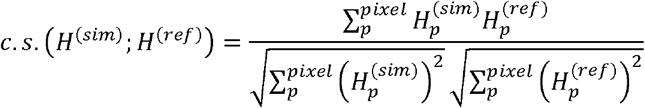

Here, *H*^(*sim*)^ denotes the pseudo-AFM image generated from a molecular model and *H*^(*ref*)^ denotes the reference AFM image. *H_p_* denotes the height at the pixel *p*. The cosine similarity takes 1.0 if two images are identical, and 0 when the two images are completely orthogonal. Notably, the cosine similarity is invariant against the uniform scaling, i.e., 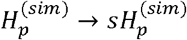. To use this function in a minimization protocol, we defined the corresponding cost function as follows.

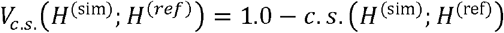

#### Correlation Coefficient

The correlation coefficient is a well-known measure that represents how two data sets resemble each other.

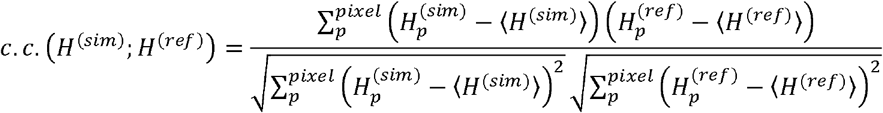

Since each term subtracts its mean value, it does not depend on the absolute height. Similar to the cosine similarity, it takes 1.0 if the two images are identical and −1.0 if images are completely opposite. Notably, the correlation coefficient is invariant against the uniform scaling, i.e., 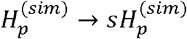, as well as the uniform shifting i.e., 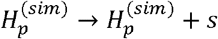. To use this in the minimization, we also defined the cost function below.

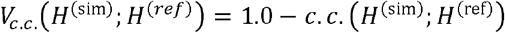

Here, 〈*x*〉 denotes the mean value of *x*’s.

#### pixel-RMSD

The root mean square displacement (RMSD) is a widely used measure to calculate the difference between two data. Here we utilized the RMSD of pixel heights as the cost function between the two images. To distinguish it from the RMSD between two structural models, we call it the pixel-RMSD hereafter. The RMSD takes 0 if the two images are identical and takes a positive value if the two images are different. The pixel-RMSD value changes by the uniform scaling and by the uniform shifting, in contrast to the above two scores. We used the pixel-RMSD value itself as the cost function in the minimization.

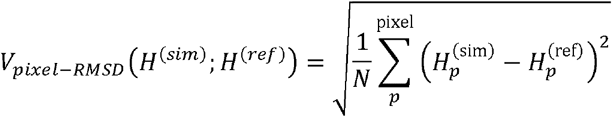

where *N* means the number of pixels.

#### Penalty function

In a previous related study, a rigid body fitting method to an AFM image using molecular docking algorithms was examined(17–19). It uses a cost function composed of three regions, forbidden, favorable, and neutral zones. Region above the topography data are the forbidden zone and particles should not be found. The region immediately beneath the topography where particle can contribute to the topography is defined as the favorable zone. The rest is defined neutral and does not affect to the cost function. The study utilized an existing docking method using the cost function, it has an advantage in terms of efficiency but, since it does not have any model of the probe, it cannot be used to reconstruct probe shape. Here, inspired by this previous approach, we define a penalty function, somewhat resemble to, but not identical to, the previous one. If a height at a pixel in a pseudo-AFM image exceeds the height of the corresponding pixel in the reference image, the cost increases. If a height of a pixel in a pseudo-AFM image is less than and close to that of the corresponding pixel in the reference image, the cost decreases. Otherwise, the cost does not change. Specifically, we define the penalty function as follows:

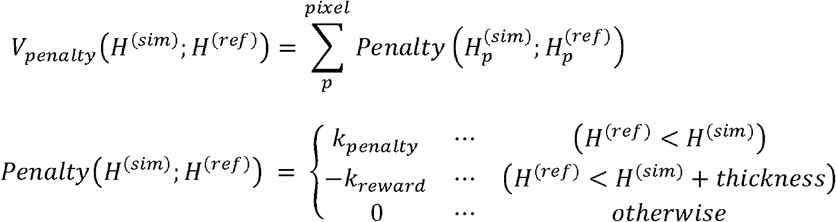

In this work, we chose 1.2 nm as the thickness of the favorable region, the same value in the previous work, 10.0 as the *k_penalty_* and 1.0 as the *k_reward_*. Note that this penalty function is not exactly the same as that of the previous research because we use pixel heights to calculate the cost but the original one uses the particle positions. Since our purpose is to reconstruct the probe shape as well as the three-dimensional structural model, we need to use a pseudo-AFM image generated with a probe model. Thus, we used heights of the pixels, not the height of the atoms in a model.

### Exhaustive Search Rigid-body Fitting Method

In this work, we exhaustively searched the discretized orientation and translation of a molecule as well as the discretized radius and the half-apex angle of the probe tip to find the optimal combination of the molecular placement and the probe shape that generates the pseudo-AFM image with the highest similarity to the reference AFM image.

To search the orientation of a molecule, we first define a step size of the rotational angle that divides 360 degree into a multiple of 4. Unless otherwise noted, we use 10 degree as the step size. Then we distribute points on a spherical surface evenly, according to the step size (Fig 1B). To distribute evenly spaced points on the spherical surface, we divided the great circle by the step size starting from the pole where the positive side of Z-axis intersects the spherical surface. Since we chose the step size to make the number of dividing points a multiple of 4, we have a point on the equator where the XY-plane intersects the spherical surface. When we call the point at the pole as the 0-th point, the 9-th point is on the equator. For each point, the n-th point, chosen on the great circle, we consider the surface that passes through the point and that is perpendicular to the Z-axis. We then divide the section of the spherical surface and the n-th plane, i.e., the nth circle, into n components. In this manner, we can distribute points on spherical surface almost evenly. Finally, we rotate the molecule to make Z-axis turns to those points, and then rotate the molecule around the new Z-axis.

For each orientation, we first generate the pseudo-AFM image and calculate the bounding rectangle of non-zero-height pixels. We then move the image translationally in XY-direction in the range where the bounding rectangle does not stick out of the reference AFM image. Here, we define the step size of the translation along the X- and Y-axis as the pixel size. We search the location of molecule along Z-axis only if the reference image contains a pixel that is higher than any of the pixel in pseudo-AFM image. Since the resolution of HS-AFM image in Z-axis is typically from 0.05 nm to 0.15 nm, we used 0.064 nm as the step size along the Z-axis.

The exhaustive search typically takes several tens of minutes on a PC with a single CPU core, although the precise time depends on the target molecules and the computer used.

### Twin Experiment

To assess the performance of the methods and compare the four similarity score functions unambiguously, we first conduct the so-called twin-experiments. A twin-experiment begins with a random choice of a position and an orientation of a target protein structure bound on a surface (see the next subsection for more details). We also choose a probe tip radius and the half-apex angle. These together serve as the ground-truth data of the molecular placement and the probe shape. Using this ground-truth setup, we generate an AFM image via the collision-detection method, to which we added 10 different realizations of Gaussian noises. The generated 10 AFM images play the role of “experimental AFM images” and are referred as the reference AFM images. Then, applying the exhaustive search rigid-body fitting method to the reference AFM image, we seek the lowest-scored placement and the probe shape.

The deviation between the ground-truth structure and the best-fit placement found in the exhaustive search is measured as the RMSD. For this purpose, we simply calculated the structure RMSD without further optimizing the alignment. To distinguish it from the pixel-RMSD used to measure the difference of two AFM images, we will call it as the structure-RMSD hereafter. Note that, since the orientation search uses relatively large step size (10 degree), the best orientation may still have non-negligible structure-RMSD.

### Protein Modeling

In this paper, we used dynein(33), myosin [pdb id: 2KIN](34), actin monomer, and actin filament [pdb id: 6BNO](35) as the targets of twin experiments. To remove the heterogeneity in the actin filament structure, we aligned the chain A in the PDB structure to the other chains and replaced them by the chain A structure. Then we aligned the last monomer of the octamer by the first monomer and obtained 15-mer and 29-mer structures of actin filament.

For the test on the real HS-AFM image of a flagellar protein FlhA_C_ ring, we used a structure of FlhA_C_ 9-mer ring obtained by placing FlhA_C_ [pdb id: 3A5I] in a ring-shaped orientation described in Terahara and colleagues (36).

In the HS-AFM experimental setup, normally proteins are weakly bound to the stage to moderate the fast Brownian diffusion. To mimic this situation while generating the reference image in the twin experiments, we calculated an oriented bounding box (OBB) of each protein and rotated the protein to ground the largest face of the OBB on the stage. The OBB axes are calculated based on the principal component analysis of the particle coordinates. The first axis is the first principal component, the second axis is the second principal component rotated to make it perpendicular to the first axis, and the third axis is the cross product of first and second axes. After generating the reference pseudo-AFM images, we rotated the model randomly around X-, Y-, and Z-axis to remove the exact reference structure from the orientation used while searching.

### Software Availability

The exhaustive search rigid-body fitting method is implemented as a part of software, the afmize(32). Originally, the afmize was a software to generate a pseudo-AFM image for a given structure, which was developed in our previous work(27). Now, it is extended to perform the rigid-body fitting. The software is freely available for download from https://github.com/ToruNiina/afmize. The method we introduced in this paper is available as the second major release. Though the time required depends on the size of the molecule, resolution of conformations, and the size of AFM image, it typically takes several tens of minutes to run on a normal desktop computer with intel i7 CPU using only one core.

### Atomic Force Microscopy Measurement and Data

For the measurement of actin filament, a laboratory-built HS-AFM apparatus built in Kodera group at Kanazawa University as described previously was used (8). Briefly, a glass sample stage (diameter, 2 mm; height, 2 mm) with a thin mica disc (1.5 mm in diameter and ~0.05 mm in thickness) glued to the top by epoxy was attached onto the top of a Z-scanner by a drop of nail polish. A freshly cleaved mica surface was treated for 5□min with 0.01% (3-aminopropyl) triethoxysilane (APTES) diluted with milli-Q water (Shin-Etsu Chemical). After rinsing the surface with drops of milli-Q water (20 μl × 5), the solution was replaced with buffer A (25 mM KCl, 2 mM MgCl2, 1 mM EGTA, 20 mM Imidazole–HCl, pH 7.6). A drop (2 μl) of actin filaments (ca. 1 μM), which were stabilized with phalloidin (11), diluted with buffer A was deposited for 10 min. After rinsing the surface with buffer A of 20 μl, the sample stage was immersed in a liquid cell filled with buffer A of 60 μl, and HS-AFM imaging was carried out in the tapping mode. We used small cantilevers (BL-AC10DS-A2, Olympus, Tokyo) whose spring constant, resonant frequency in water, and quality factor in water were ~0.1 N/m, ~0.5 MHz, and ~1.5, respectively. The probe tip was grown on the original tip end of a cantilever through electron beam deposition using ferrocene and was further sharpened using a radio frequency plasma etcher (Tergeo, PIE Scientific LLC., USA) under an argon gas atmosphere (Direct mode, 10 sccm and 20 W for 1.5 min). The cantilever’s free oscillation amplitude A0 and set-point amplitude As were set at ~2 nm and ~0.9 × A0, respectively. Details of the method for HS-AFM imaging are described elsewhere (37).

For the AFM movie of FlhA_C_, we used the data obtained previously (36), in which the experimental parameters relevant to the current study are the followings: A glass sample stage (diameter, 2nm; height, 2nm) with a thin mica disc (1mm in diameter and ~0.05 mm thick) glued to the top by epoxy was attached on to the top of a Z-scanner by a drop of nail polish. A fleshly cleaved mica surface was prepared by removing the top layers of mica using a Scotch tape. AFM imaging was carried out in a tapping mode, using small cantilevers (BLAC10DS-A2, Olympus).

## Results

### Noise analysis in experimental AFM images

Before the setup of twin-experiments, we analyzed several AFM images to estimate the intensity of the background noise in the observation. We took two HS-AFM movies that observe FlhA_C_ monomers(36) and actin filaments because those movies contain large background areas and selected 5 frames in the last part of the movies. Then we manually selected background regions from those images and estimated the location of the stage via the standard least square method assuming that the stage is a tilted plane. After that, we collected the deviation of the height of each pixel from the estimated stage position.

Fig S1 shows the histogram of the deviation collected from the background area in the experimental observation of FlhA_C_ monomers (the first case) and actin filaments (the second case). In the first case, the histogram fits well to the Gaussian of which the standard deviation is 0.284 nm. In the second case, although the distribution slightly deviates from the Gaussian because of some outliers, we fitted it with a Gaussian of which the standard deviation is 0.347 nm. In the second case, we recognized several unidentified objects that are supposed to be contaminants, which could lead to a small deviation of the stage position and cause the distortion of the histogram. Since this appears only in limited area, this is not taken into accounts here. We concluded the noise in the HS-AFM observation can be approximated by the normal distribution with its standard deviation around 0.3 nm. We used this value as a noise level when we generate the reference pseudo-AFM image in the twin experiments.

### Twin-experiment: Dynein

As the first test protein, we chose dynein because of its large size and anisotropic shape. Since the random noise in the AFM image is independent from molecules, the larger the target molecule is, the higher signal-to-noise ratio we will obtain. Also, its anisotropic structure is expected to make the orientation search relatively easy.

We placed a dynein model(33) on a surface and generated two reference-AFM images using two different probe shapes, 1 nm / 10 degree (Fig 2A) and 3 nm / 20 degree (Fig 2C) (for the probe shape, the values before and after the slash indicate the tip radius and the half-apex angle, respectively, throughout). These two images are treated as the reference “experimental” AFM images in the context of the twin-experiment. For each reference-AFM image, we applied the exhaustive search method using the four different score functions and compared all the results. Since the probe tip radius used in real AFM imaging(8) is normally between 0.5 nm and 5 nm, we used tip radii of the same range in the search. After obtaining the minimum cost structure, to evaluate the accuracy of the structure placement we calculated the structure-RMSD with respect to the ground-truth structure used to generate the reference AFM image. Note that the exhaustive search method uses the discrete space for the orientation (10 degree here), the translation (same as the pixel size. typically, it is 1 nm), the tip radius (6 values from 0.5 nm to 5.0 nm), and the half-apex angle (10 and 20 degrees). By applying random translation/rotation to prepare the initial structure, we did not include the exactly-correct position/orientation in the test cases. Thus, the minimum possible structure-RMSD may become the same order of sum of the half step sizes.

**Fig 2.**
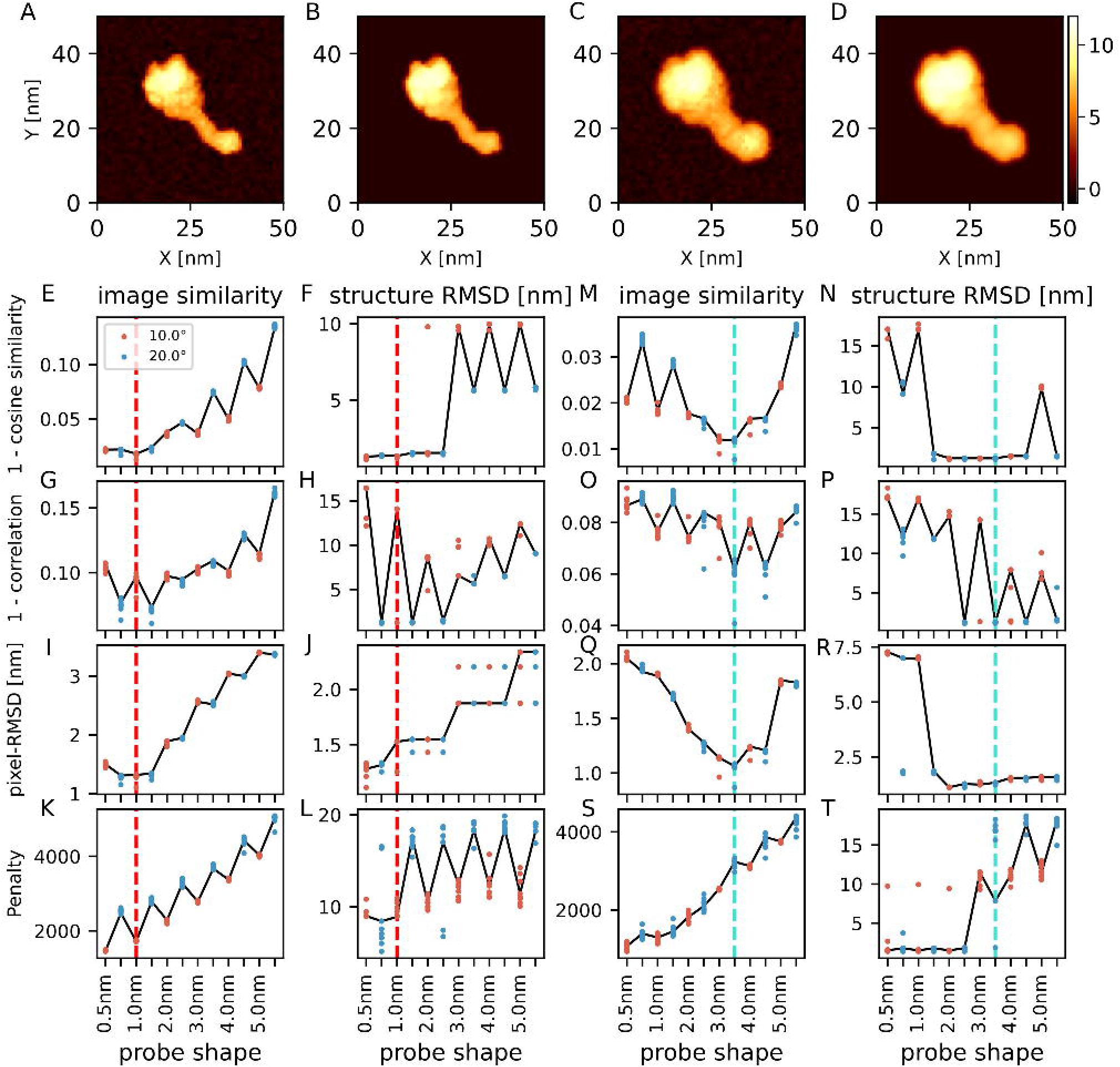
Twin-experiments for a molecular model of dynein. (A) One of the reference AFM images in the twin-experiment, with the 1 nm / 10 degree probe. (B) The image generated from the predicted structure with the cosine similarity. (C) One of the reference AFM images with the 3 nm / 20 degree probe. (D) The image generated from the predicted structure with the cosine similarity. The color bar shows the height in nm (shared by all of the images here). (E-L) The results of twin-experiment with the reference image generated by the 1 nm / 10 degree probe. Results of 10 replicated runs are overlaid; each run uses different reference AFM images with independent noise. The row corresponds to the cost function. The leftmost 4 panels (E, G, I, K) show the lowest scores of images. The next 4 panels (F, H, J, L) show the structure-RMSD of the structures with the lowest score. Red vertical dashed lines show the ground-truth probe shape. Results from two probe angles, 10 degree in red and 20 degree in blue, are plotted in parallel. Solid lines connect the representative results from one reference AFM image. (M-T) The results of twin-experiment with the reference image generated by 3 nm / 20 degree probe, shown in the same way as E-L. The cyan vertical dashed lines show the ground-truth probe shape.

Fig 2E-L show the result of the case when the reference-AFM image is generated with the 1 nm / 10 degree probe. When we used the cosine similarity as the cost function, the lowest cost configuration across all the search space was always the one with correct probe shape. The structure-RMSD of those configurations were around 1.3 nm and it is smaller than that of the structures predicted by different probe shapes (5.6 or 9.8 nm). Contrary, to our surprise, when we used the correlation coefficient, the lowest cost configuration was not the correct probe shape but the probe with 1.0 nm / 20 degree. In the resulting structure, the dynein molecule is placed vertically to fit to the AAA ring. This might be caused by two reasons, masking and the scale invariance to the uniform shift of the correlation coefficient. As described in Methods, unlike the cosine similarity, the correlation coefficient has the same value after adding a constant value across an image. Since pixels that have zero height in the pseudo-AFM image are ignored in the cost function in our approach, the correlation coefficient sometimes falls into an overfitting to some portion of the image. The structure-RMSD of those configurations were almost the same as that of the cosine similarity (~1.3 nm). Thus, it can reconstruct the orientation of molecule even though it cannot accurately infer the probe tip angle. When we used the pixel-RMSD as the cost function, the lowest cost configuration was rarely in the correct probe shape (3 out of 10), but was 0.5 nm / 20 degree (7 out of 10). However, the structure-RMSD of the lowest cost configuration searched with a different probe shape was comparatively small compared to other cost functions (~1.3 nm). Lastly, when we used the penalty function as the cost function, the result with the smallest probe shape had the lowest cost value. This is reasonable because the penalty function increases its costs if a pixel in a pseudo-AFM image exceeds that of the reference AFM-image, and smaller probe shapes tend to produce images that contain less non-zero pixels because of the small cross section and also make the pixels around the concave regions of molecular surface lower.

Fig 2M-T show the result of the case when the reference-AFM image is generated with the 3 nm radius / 20 degree probe. In this case, the larger tip radius tends to hide fine features on the surface of the specimen, so the structure placement must be more difficult. When we used the cosine similarity as the cost function, the lowest cost configuration was often reached with the correct probe shape (i.e., 3 nm radius / 20 degree)(7 out of 10), but sometimes the 3 nm radius / 10 degree probe had the lowest cost (3 out of 10). The structure-RMSD of those configurations were as small as that in the case of 1nm radius / 20 degree probe (~1.3 nm). When we used the correlation coefficient as the cost function, the results of the correct probe shape often had the lowest cost (7 out of 10) and the structure-RMSD was also low (~1.3 nm). In the case of the pixel-RMSD, the results of the correct probe shape always had the lowest cost and the structure-RMSD was low (~1.3nm). The penalty function favors, as before, the smallest probe.

In summary of the dynein case, the cosine similarity generally worked well and, in the case of relatively blunt probe, the pixel-RMSD and the correlation coefficient worked well, too.

### Twin-experiment: Myosin

Next, we conducted otherwise the same examination for a myosin motor domain complex. The myosin motor domain is smaller in size than dynein so that the structure placement is expected to be more difficult than the case of dynein. The results for myosin showed overall the same tendency as the case of dynein (Fig S2). Just in summary, the cosine similarity score worked well both in the structure placement and the probe shape inference in most cases. The correlation coefficient worked well in the structure placement, but it sometimes failed to infer the probe shape, maybe because of the same reason described in the case of dynein. The pixel-RMSD worked better when the reference-AFM image is generated by a larger probe, but otherwise, worked slightly worse. The penalty function always favors smaller probe shape and the structure-RMSD of best-score structure was small.

### Twin-experiment: Actin monomer

We further applied the same method to an actin monomer, which is even smaller than the myosin motor domain, and thus is a harder target.

Fig 3E-L show the result when the reference-AFM image is generated by the 1 nm / 10 degree probe. When the cosine similarity is used as the cost function, it had less probability to infer the correct probe shape (4 out of 10) and it was difficult to distinguish the correct one from similar probes, such as 0.5 nm / 10 degree (5 out of 10). The placed structure sometimes had markedly large structure-RMSDs, such as 3 nm, where the orientation was almost opposite. We note that the actin monomer is only weakly anisotropic so that it is rather difficult to distinguish the correct orientation from the opposite one. The prediction by the correlation coefficient showed nearly the same tendency as the cosine similarity. When we used the pixel-RMSD as the cost function, the reconstructed probe shape is rather different from the correct one, but, to our surprise, the placed structure always had relatively small structure-RMSDs. This is possibly because the pixel-RMSD directly compares the height of the image without the invariance to the uniform scaling and can decide the placement more robustly in the idealized situation of the twin-experiment. The penalty function showed the same severe artefact as the cases of dynein and myosin.

**Fig 3.**
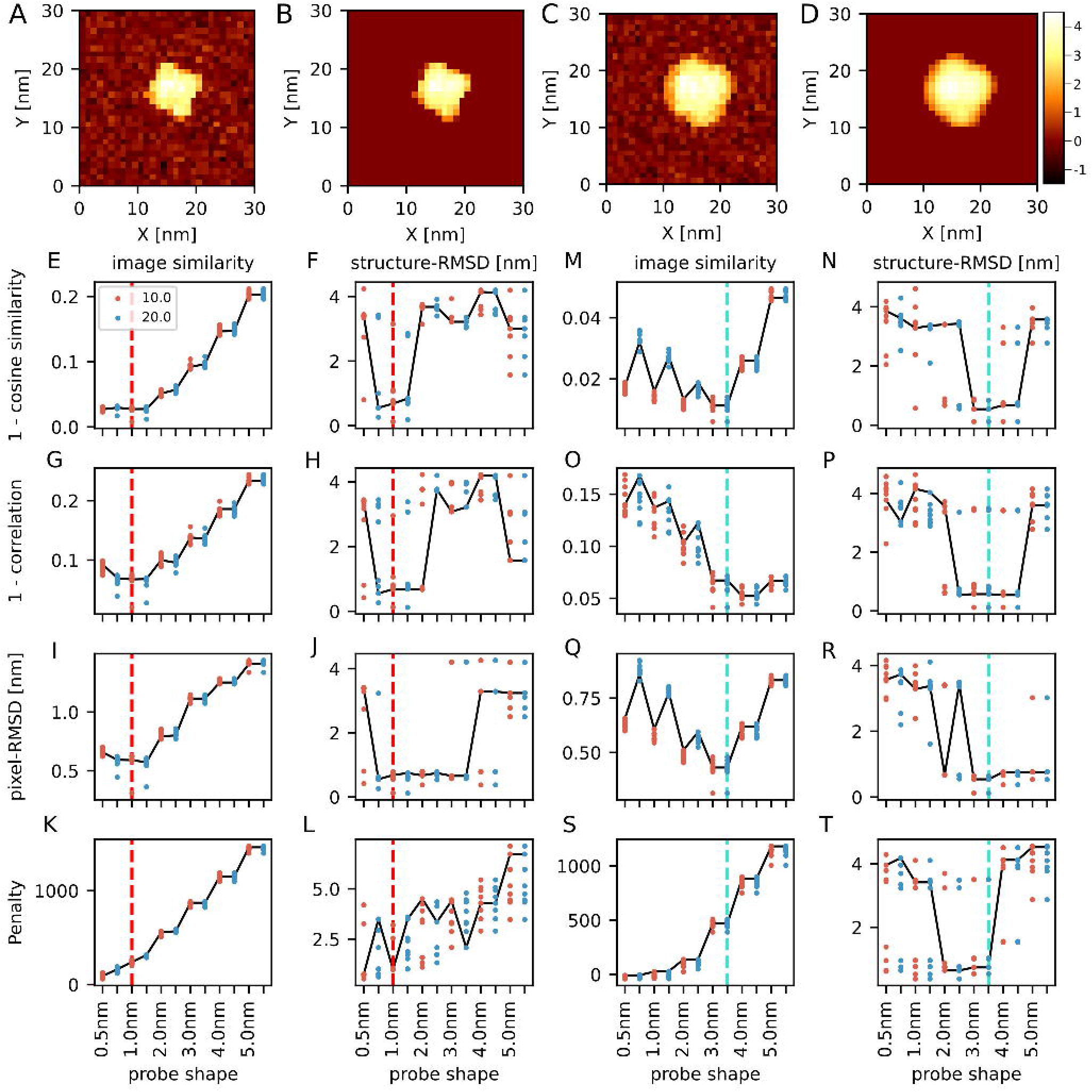
Twin-experiments for an actin monomer. (A) One of the reference images in the twin-experiment, generated with the 1 nm / 10 degree probe. (B) The image generated from the predicted structure with the cosine similarity-based cost function. (C) One of the reference images in the twin-experiment with the 3 nm / 20 degree probe. (D) The image generated from the predicted structure with the cosine similarity-based cost function. The color bar shows the height in nm (shared by all of the images here). (E-L) The results of twin-experiment with a reference image generated by the 1 nm / 10 degree probe. Results of 10 replicated runs are overlaid; each run uses different reference AFM images with independent noise. The row corresponds to the cost function. The leftmost 4 panels (E, G, I, K) show the lowest scores of images. The next 4 panels (F, H, J, L) show the structure-RMSD of the structures with the lowest score. Red vertical dashed lines show the ground-truth probe shape. Results from two probe angles, 10 degree in red and 20 degree in blue, are plotted in parallel. Solid lines connect the representative results from one reference AFM image. (M-T) The results of twin-experiment with the reference image generated by the 3 nm / 20 degree probe, shown in the same way as E-L. The cyan vertical dashed lines show the ground-truth probe shape.

Fig 3M-T show the result when the reference AFM image is generated by the 3 nm / 20 degree probe. Using the cosine similarity, it could not infer the probe shape. The structure-RMSD of the best fit configuration sometimes had a large value. The correlation coefficient showed almost the same as the case of the cosine similarity. However, when the pixel-RMSD was used as the cost function, the reconstructed configuration always had relatively small structure-RMSD. The penalty function favors too small probe, as before. All the cost functions did not distinguish the tip apex angle when the probe radius exceeds 3 nm. This is because the highest pixel in the pseudo-AFM image has only 4 nm, thus the conical part of the tip almost never interacts with the protein when the probe radius is larger than 3 nm and the resulting pseudo-AFM image becomes the same.

From those results, when the target protein is relatively small and the target image has pseudosymmetry, we found that the pixel-RMSD works better than the cosine similarity and the correlation coefficient in terms of the prediction of the placement. When the probe shape is acute, the pixel-RMSD could infer the probe shape, but the accuracy decreases. So, in this case, it could be better to use the pixel-RMSD to find the best configuration and later compare the probe shape by using the cosine similarity. Note that, the pixel-RMSD highly depends on the absolute value of the heights. Thus, when applying the pixel-RMSD to the experimental data, we need to estimate the stage position in Z direction accurately.

### Twin-experiment: Actin filament

As the final case of twin-experiments, we examined a case where we fit a partial molecular structure into a reference AFM image that contains a larger molecular complex. We generated a reference AFM image using 29-mer structure of actin filament (the AFM image in Fig 4AC), which serves as the ground-truth. Note that the actin filament has the directionality; the one end is called the barbed (+) end, and the other called the pointed (-) end. We then used a 15-mer of actin as the structure to be fitted (Fig 4BD). To calculate the structure-RMSD between the predicted structure and the ground-truth structure, we calculated the structure-RMSD between all possible consecutive 15-mer in the reference 29-mer structure and took the minimum value as the structure-RMSD between the prediction and the ground-truth structures.

**Fig 4.**
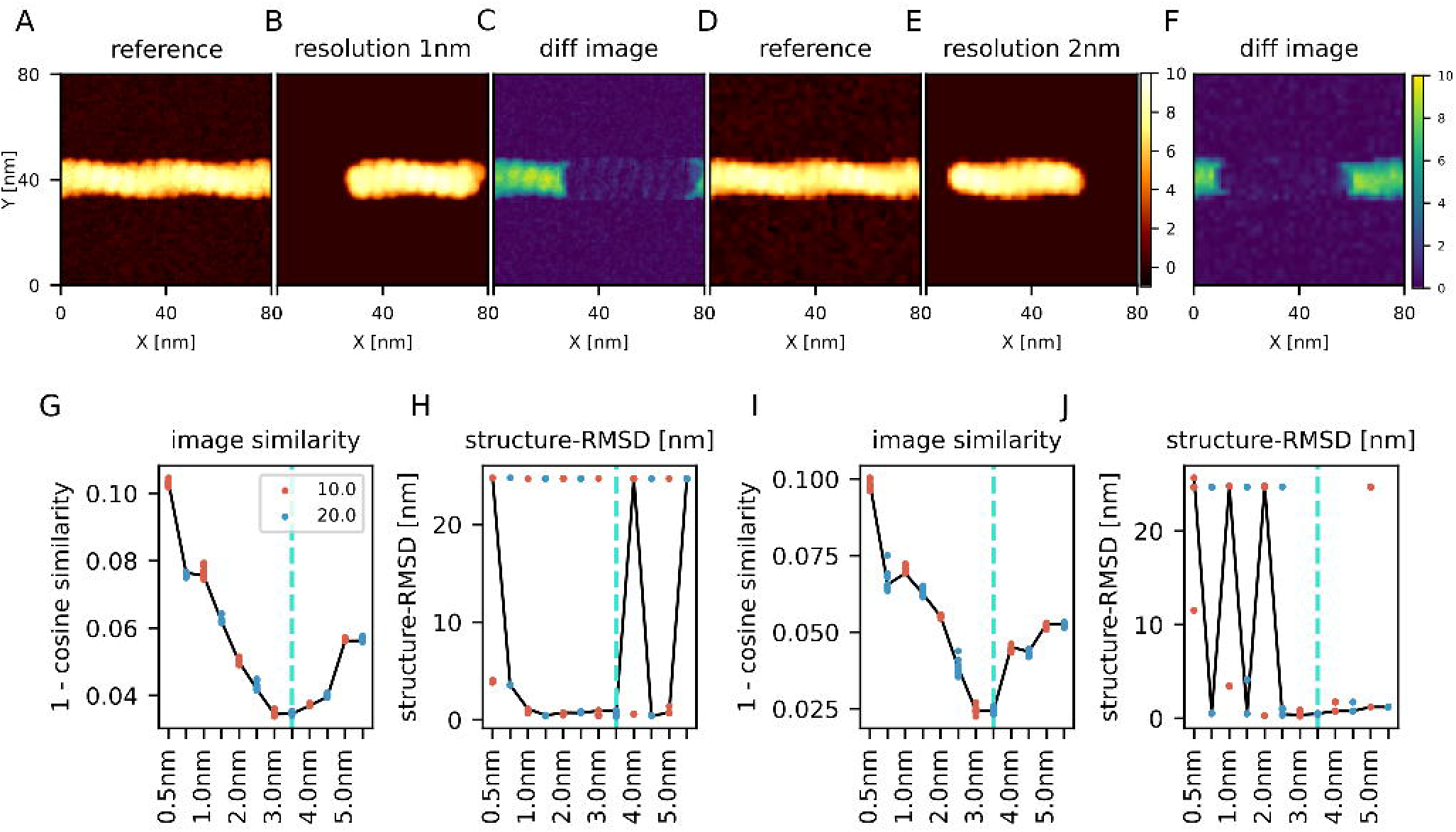
Twin-experiment for actin filament. (A) One of the reference images used in this twin experiment, generated by a 3 nm / 20 degree probe, with the pixel width 1 nm. (B) The image generated using the structure predicted from the image (A) with the cosine similarity-based cost function. (C) The difference between (A) and (B). (D) One of the reference images used in this twin experiment, generated by a 3 nm / 20 degree probe. Unlike the panel (A), the pixel width is 2 nm, which is twice as large as that of panel (A). (E) The image generated from the structure predicted from the image (D) with the cosine similarity-based cost function. (F) The difference between panel (D) and (E). (A,B,D,E) The colormap of all the pseudo-AFM images is shared. (C,F) The color-bar of the difference maps is shared. (G) The resulting best scores of the prediction from the image with 1 nm pixel. (H) The structure-RMSD between the ground-truth structure and the predicted structure from the image with 1nm pixel. (I) The resulting best scores of the prediction from the image with 2 nm-wide pixels. (J) The structure-RMSD between the ground-truth structure and the predicted structure from the image with 2 nm-wide pixels. (G-J) Results from two probe angles, 10 degree in red and 20 degree in blue, are plotted in parallel. Solid lines connect the representative results from one reference AFM image. The cyan line shows the ground-truth probe shape.

First, same as before, we set the pixel/grid size to 1 nm and performed the twin-experiment using the cosine similarity with the same set of probe shapes. The method found the correct probe shape as the lowest cost (8 out of 10) and the structure-RMSD was well small in those configurations (Fig 4EF). Fig S3 shows the result with the other three cost functions. Except the penalty function, other cost functions also found the correct probe shape and the configuration rather accurately. From those results, both the probe shape and configuration can be estimated by using a portion of the complex structure.

Second, we performed the same twin experiment with the image resolution as the pixel/grid size as 2 nm (Fig 4GH), which is twice as large as the previous case. Using the cosine similarity, the probe shape can often be estimated correctly (7 out of 10), or close to the correct shape (3 nm radius /10 degree) otherwise. The best-fit placements had small structure-RMSDs.

Fig S3 shows the result with the other cost functions. Use of the correlation function correctly estimates the probe radius, but fails to infer the probe angle. With the pixel-RMSD, it often finds the correct probe shape (6 out of 10), but it sometimes has incorrect apex angle of 10 degree (4 out of 10). We found that sometimes the resulting structures have large structure-RMSD. In these cases, the location of the structure was fine, but the directionality of actin filament was opposite. This failure suggests that, in case of near-symmetric target molecules with a low resolution of AFM image, the structure inference could be slightly difficult.

From those results, we found that the cosine similarity, the correlation coefficient, and the pixel-RMSD work well for both the probe shape inference and the structure placement when we have an image with 1 nm resolution. However, we also show that when we have an image with the coarser resolution, 2 nm per pixel, the cosine similarity and the pixel-RMSD worked relatively better and the other cost functions could estimate probe radius but not the apex angle of the probe.

### Real-data: Actin filament

So far, we applied the exhaustive search rigid-body fitting method to the synthetic-AFM images, i.e., the twin-experiments because we need the ground-truth structure for assessing the accuracy of the predictions. Now, we apply our method to real experimental AFM images. Here we used an HS-AFM image of the actin filament on which we fit the 15-mer structure of actin filament.

First, before the application, we estimated the stage surface. The stage surface is close to, but, inevitably, not perfectly parallel to, the *XY*-plane of the AFM apparatus. Thus, the measured Z-coordinate does not directly represent the height of the specimen. For accurate modeling, we first estimated the stage surface and corrected the measured data. To estimate the stage position from the experimental image (Fig 5A), we extracted pixels that are supposed to represent the stage and fitted these pixels by the plane surface via the least square method (Fig 5B). The obtained equation of the surface plane was *z* = 0.0004*x* – 0.003*y* – 18.8 (*nm*). The surface is indeed close to, but not identical to the *XY*-plane; it has almost 1 nm difference between the edge of the image (Fig S4). We used this equation and subtracted the height of the stage from the apparent data (Fig 5C) at each pixel and chose 40×40 pixels region as a reference AFM image from the corrected image (Fig 5D).

**Fig 5.**
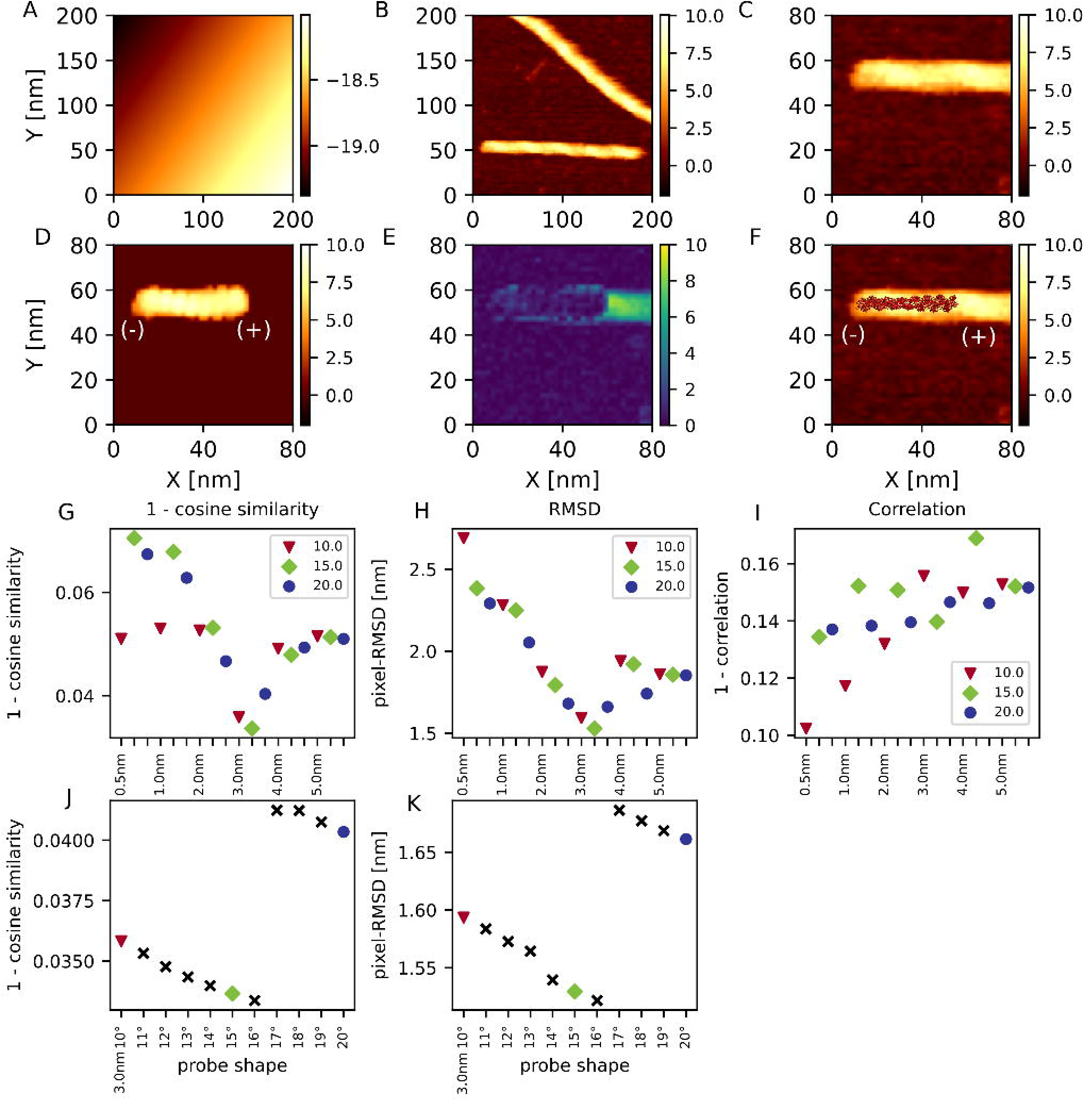
Exhaustive search rigid body fitting to an experimental HS-AFM image of actin filament. (A) The stage plane estimated from the background regions of experimental AFM image. (B) The AFM image after the correction of stage position. (C) The reference image used in the rigid-body fitting. It is taken from the lower left rectangular region of the image in B. (D) The pseudo-AFM image generated by the best estimation with the cosine similarity score with the 3 nm / 16 degree probe that shows the best score. (E) The absolute value of difference between C and D. (F) The resulting actin filament structure model on top of the reference AFM image. In (A-F), the color map is given in nm unit. In (D) and (F), the orientation of the generated actin filament model is indicated by (+) and (-) labels. (G-I) The cost values of the best fit using varying probe shape. From left to right, the result using the cosine similarity (G), the pixel-RMSD (H), and the correlation coefficient (I) are plotted. In (G-I), the marker color and shape represent the probe angle and three consecutive markers have the same probe radius. (J, K) The results of finer-grid search of probe angle with radius 3.0, using the cosine similarity score (J) and the pixel-RMSD score (K).

For the corrected HS-AFM image given in Fig. 5BC, Fig 5EF show the result of the exhaustive search rigid-body fitting with the cosine similarity. The probe shape that achieves the lowest cost was 3.0 nm / 15 degree (Fig 5G). Therefore, the actual probe shape seems to be around this value. The result of pixel-RMSD was consistent with these values (Fig 5H). This value is in the range of the typical AFM probe radius 0.5 nm ~ 5.0 nm in literature (8). Then, we further searched the probe shape at a finer grid between 10 degrees and 20 degrees with 3 nm radius (Fig 5JK). Both the cosine similarity and the pixel RMSD gave the best score with the probe 3 nm / 16 degree. This consistence shows the reliability of the result.

The result with the correlation coefficient was not consistent with the other two cost functions. In the best fit result with the correlation coefficient, the actin filament was placed almost vertically to fit the left terminal of the filament. This ill-behavior is similar to that found in the case of twin-experiment for dynein.

Finally, we address the inference/prediction of the orientation of the actin filament in the AFM image. We first note that, we cannot identify the orientation of the filament by eye. It should be noted that, experimentally, the actin filament orientation can be investigated by the binding of some actin-binding proteins. In the previous section, the twin-experiment showed that our method can identify the orientation accurately for the image in the ideal condition. Then, what about the case of a real AFM image? In the top 10 alignment with the 3.0 nm / 16 degree probe, we found all the top 10 structures having the barbed (+) end points at the right. The pixel-RMSD function showed the same results. We note that, in the all top 10 alignment, the structure is placed at the edge of the filament of the AFM image. Thus, the presence of the edge in the image could help identifying the orientation. In summary, the orientation of the actin filament is likely to be identified by the exhaustive-search rigid-body fitting, even when it is hardly recognized by visual inspection.

### Real-data: FlhA_C_ ring

Lastly, we applied the exhaustive search rigid-body fitting to the ring-like FlhA_C_ homo oligomer(36) (Fig 6). To estimate the stage position, we first extracted several small regions that are supposed to represent the stage. We performed the least square fitting to the equation of a plane (Fig S5AB), and subtracted the height of the stage (Fig 6AB). As a target image, we then selected a ring at the left side of the image and extracted a rectangular region where the ring is visualized (Fig 6C). We used a ring shaped nonamer structure for the rigid-body fitting with the cosine similarity, the pixel-RMSD, and the correlation coefficient as the cost function.

**Fig 6.**
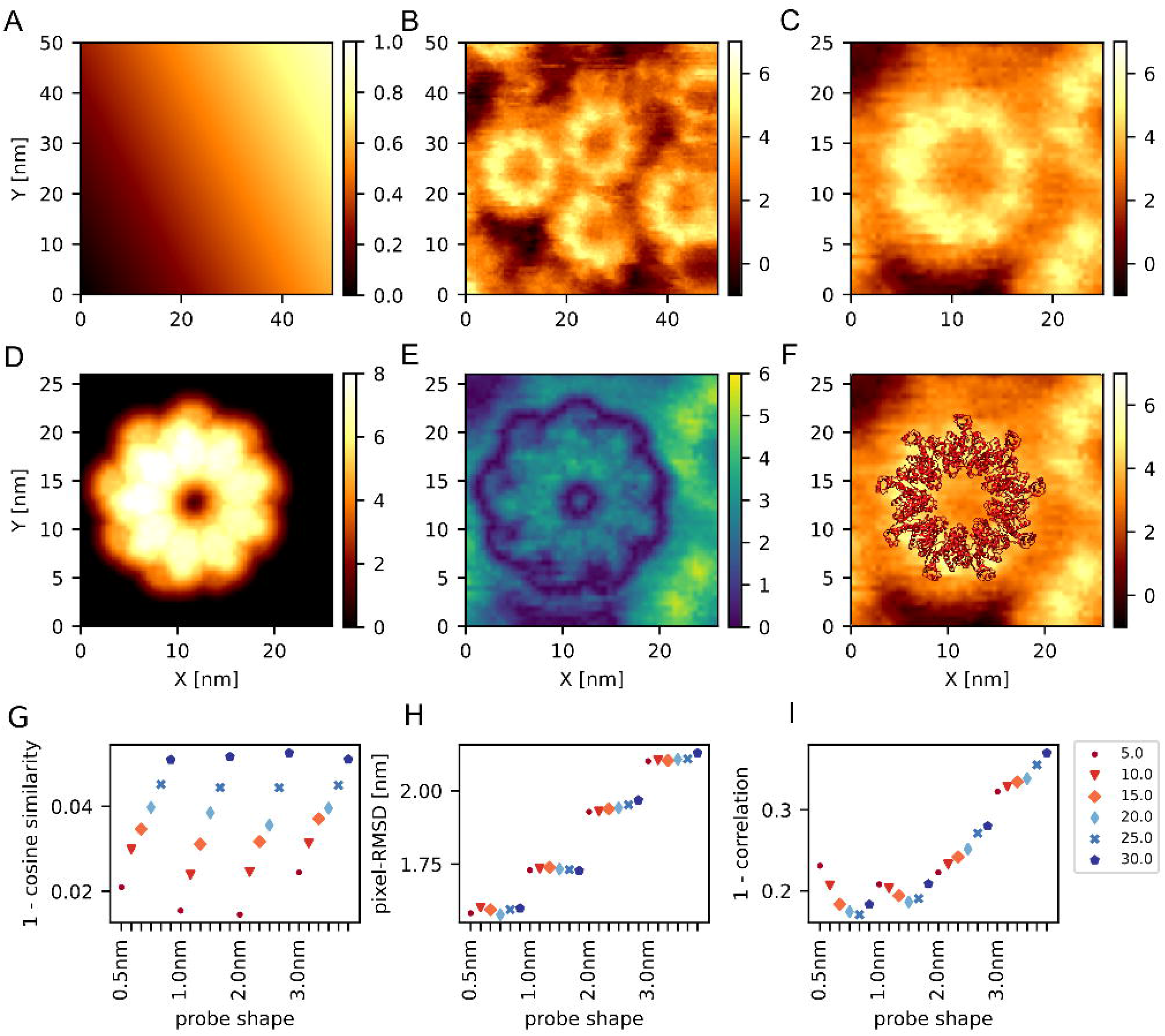
Exhaustive search rigid body fitting to an HS-AFM experimental image of FlhA_C_ ring (36). (A) The stage plane estimated from the background regions of experimental AFM image. (B) The AFM image after the correction of stage plane. (C) The reference image used in the rigid-body fitting. It is taken from the left middle rectangular region of the image in (B). (D) The pseudo-AFM image generated by the estimation result with the correlation coefficient. (E) The absolute value of differences between (C) and (D). (F)The resulting FlhA_C_ nonamer ring structure model on top of the reference AFM image. (G-I) The result of fitting using cosine similarity (G), pixel-RMSD (H), and correlation coefficient (I). The colors and shapes of the marker represents the angle of the probe. The angle is 5 (dot), 10 (triangle), 15 (diamond), 20 (thin diamond), 25 (x), and 30 degrees (hexagon), respectively. Within 6 trials using varying probe angles, the same probe radius is used.

In any cases, we found that the resulting pseudo-AFM images generated from the best-fit have markedly larger height compared to the experimental AFM image by ~1.5 nm (Fig 6DE for the case of the correlation coefficient). Looking into the structure model of FlhAc, we noticed that the molecule has relatively long loop region at the top and bottom in the ring structure, which is in contrast to the structure of the actin filament that does not have long loops. Obviously, such loop structures can easily be altered by the interactions with the stage and/or the tip. The rigid-body fitting, in its current form, does not account for such plasticity at all. Thus, the rigid-body fitting inevitably gave markedly higher pseudo-AFM images than the real AFM image.

If we assume that the loop structure is largely compressed whereas the other core domain keeps its reference structure in the AFM measurement environment, the invariance of the similarity score with respect to the uniform shift i.e., 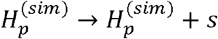, becomes a key property, where *s* corresponds to the compression of the loop. The uniform scaling could also account for some of structure deformation. Among the three similarity scores, only the correlation coefficient has the invariance with respect to the uniform shift and the uniform scaling. Actually, when we apply a linear scaling and an uniform shifting to the best-fit structure with the correlation coefficient, the transformed image becomes much closer to the real AFM image (Fig S5 D-J). It could represent the compression in the loop region on top of globular part. Notably, if we restrict shifting to negative z direction, shifting does not help to decrease the pixel-RMSD and the result was the same as the case in which we applied scaling only. In contrast to the correlation coefficient, both the pixel-RMSD and the cosine similarity are sensitive to the uniform shift of the image. In fact, visual inspection of the best-fit images clearly suggests that the image obtained by the correlation coefficient is superior to those by the others (Fig S5 M-P).

Not surprisingly, the inferred probe radius and angle were rather different among the three (Fig 6G-I); none of the pairs were consistent. The correlation coefficient exhibited a minimum at 0.5 nm radius and 25.0 degree.

## Discussion

In this work, we developed the exhaustive search rigid-body fitting method to infer the placement and the probe tip shape that generate most similar pseudo-AFM image to the experimental/reference AFM image. Although there are some limitations, by applying the method with varying probe shape parameters, we showed that the shape of the probe used in the reference AFM image can be inferred. By comparing four score functions in the ideal twin-experiments, we showed that the cosine similarity score generally works well and both the pixel-RMSD and the correlation coefficient are also useful. We also applied the rigid-body fitting to the two real HS-AFM images, the actin filament and the FlhA_C_ ring, and estimated the shapes of the probes, finding that the appropriate similarity score differs between the two cases. The cosine similarity was the best in the actin filament image, while the correlation coefficient was the best in the case of FlhA_C_ ring. The difference can be attributed to the plasticity of the target molecules. Which similarity score is best for a given target is not *a priori* clear so that, in practice, we recommend users to apply the method with all the three similarity scores. Probably, one can pick up the best score and its best molecular placement via visual inspection. Since the shape of the probe strongly affects the AFM image, the probe shape will be very important information for further refinement analysis, such as the flexible fitting.

By comparing the four score functions via twin experiments of the grid search method, we showed that the cosine similarity works well under many conditions like different image resolution, tip shape, and the sizes of molecules, both in the probe size estimation and the configuration estimation. The cosine similarity is a function that resembles to the correlation function and has been extensively used in cryo-EM flexible fitting to compare the electron density map(28) and also in the AFM flexible fitting to compare the reference AFM images(27). In the case of small proteins, the pixel-RMSD can work better than the cosine similarity in terms of the placement inference. Therefore, in the cases of relatively small proteins, it might improve the estimation to use the pixel-RMSD to determine the location and the orientation of the protein of interest. After that, the probe shape can be improved by comparing values from cosine similarity cost functions using the pseudo-AFM images generated with the structure obtained by pixel-RMSD calculation.

In fact, these two scores, the cosine similarity and the pixel-RMSD, are closely related mathematically. To understand the relation, we introduce the pixel heights of both reference and pseudo-AFM images normalized to unity, 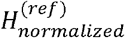 and 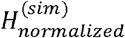 respectively. Then, the pixel-RMSD and the cosine similarity are related by

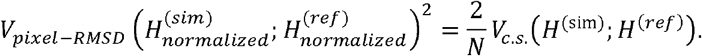

Namely, the score for the cosine similarity of the original (*unnormalized*) images is proportional to the square of the pixel-RMSD of the *normalized* images. Therefore, only the presence or absence of the normalization is the source of the difference between the two scores. Generally, two images generated with two different probe radii have noticeably different heights only at clefts and cavities on the molecular surface, and the edges of the molecule. This results in relatively small pixel-RMSD values of *unnormalized* images with different probe radii. On the other hand, the sum of the squared heights, which appeared in the normalization factor, depends on the probe radius; it is larger with a larger probe radius. Thus, when the pixel heights of the two images are normalized with different probe radii, the heights would differ at all the pixels, due to the different normalization factor, resulting in a larger pixel-RMSD of the *normalized* images, i.e., the score for the cosine similarity. This property may increase the pixel-RMSD of *normalized* images with different probe radii and contribute to the improved ability of the cosine similarity to detect the difference in probe radii.

In the application to the real AFM image of the FlhA_C_ ring, we found the correlation coefficient is the best score among the three scores examined. This superiority of the correlation coefficient for the FlhA_C_ ring can be attributed to its invariance of the score with respect to the uniform shift and the uniform scaling. When the target molecule contains noticeable loops and other flexibility, these two types of invariance would help the rigid-body fitting. It should be noted that the correlation coefficient, in our current implementation, can fail to estimate the molecular placement in some cases (the case of the twin-experiment of dynein and the real AFM image of the actin filament). However, the resulting molecular placement often has obviously different height distribution from the target image. Thus, these failure can be avoided just by supervising the results by the user.

We found that the penalty-based cost function is not applicable to the probe size estimation because it always favors smaller probe that produces smaller image. This is because if there are less non-zero pixels in the model AFM image, the opportunity to have a penalty decrease. Because of this feature, sometimes it fails to estimate the configuration. We note that, this penalty-based function is not the same as that used in the previous research. The penalty-based cost function used in this work calculates cost using the height of pixels in an AFM image, and the previous research calculates the cost using coordinates of particles.

We here used a rigid-body collision detection method to generate pseudo-AFM images, but in reality, there might be some deformation of biomolecules under the AFM measurement. To go beyond this limitation, we could estimate the elasticity of the protein to estimate a scale factor on the heights in the pixel using some other methods, such as molecular dynamics simulations, although these methods are much more time-consuming.

In both of two experimental AFM images, the estimated probe angle was extremely sharp (5 degree or less), compared to the value guessed in other researches(25,26), which were around 20 degree. It is possible that the apex-angle differs largely sample by sample. Alternatively, there could be another possibility that the observation method of HS-AFM, the tapping mode in the HS-AFM, caused this difference. Since, in the tapping mode, the probe always vibrates and only taps the sample and quickly detaches from it, the lateral part of the probe has less chance to collides with the sample. Thus, the resulting image could contain the effect of tapping and the estimated probe shape might be affected by those artifacts. Although the estimated probe shape is possibly affected by the observation technique, the result of estimation can be used as an effective probe shape in the successive analysis, such as flexible fitting, because the systematic error caused by the effective tip radius can be described by the current tip model. To decipher these points, it may be interesting to include the vibrating cantilever and probe tip controlled via a feedback within the molecular simulation systems, although this makes the estimate much more time consuming.

In this paper, for simplicity, we modeled a probe tip as a combination of simple geometric shapes that can be controlled only by two parameters, the hemisphere radius and the half apex angle of a circular frustum of a cone. In reality, however, the probe shape could have much more complicated shape. There is a classic method to estimate the probe shape from an AFM image without any prior knowledge, called the blind tip reconstruction method (39–41). Since the method does not postulates the shape of the probe, this is much more ambitious and thus a potentially more powerful method. Notably, the blind tip reconstruction has not been used in biomolecular AFM experiments partly because the method is highly sensitive to noise (25). In the present study, we briefly tested the applicability of the blind tip reconstruction algorithm for biomolecular AFM, using the twin-experiment sample and the real HS-AFM image of the actin filament. In the twin-experiment of the actin filament, we first used the noise-free pseudo-AFM image from our structure model (Fig S6A) and applied the blind tip reconstruction method (Fig S6BC). The results suggest a successful estimate in the cross-section along *Y*-axis, but not so along *X*-axis. Second, when we added the standard noise that is the same level as the above studies, the inferred tip shape becomes highly sensitive to the noise (Fig S6D-H). Next, for a real HS-AFM image of the actin filament (Fig S7A), we applied the blind tip reconstruction method, obtaining rather blurred shapes estimated. In particular, the radius of the reconstructed apex sensitively depends on the unknown threshold parameter in the method (Fig S7B-E). Overall, we conclude that the blind tip reconstruction, albeit potentially ideal, cannot easily be applied to biomolecular AFM images.

Technically, searching the best location of a molecular model can be speeded up by using the fast Fourier transformation (FFT)(38). However, within our program, we found that the most time-consuming operation was not the translational search, but the pseudo-AFM image generation for all the discrete orientations of molecules. Also, as the FFT imposes periodicity and wraps around the borders of AFM images instead of just cropping, it could lead to incorrect score values when the molecule(s) located at the borders of the image. Therefore, we did not use the FFT-based method here for the translational search. In the AFM image case, the data are two-dimensional images and the grid size (~1nm) is larger than that of the docking simulation (~0.1 nm) where the FFT is of more importance. We have much less grids (typically, the order of 10^3^ or 10^4^ after focusing on a region of interest) compared to the three-dimensional docking simulation (typically, 10^6^ grids).

In our development stage, we tried to use the Monte Carlo simulated annealing for the search process, as in our previous study(27). The Monte Carlo process includes translational and rotational movements of the target molecule, as well as the change in probe tip shape. This stochastic search worked well in some cases when the temperature cooling schedule was fine-tuned. We, however, found that the temperature cooling schedule that works well differs case by case and thus must be carefully tuned for each target system. Often, this tuning was not straightforward. In addition, we need to repeat many independent runs to confirm that we reached the global minimum, without being trapped to a metastable basin. Altogether, we concluded that the simulated annealing is not the best approach for this rigid-body fitting, and decided to employ the exhaustive search protocol, which does not need any simulation parameters to be tuned and is more robust. We anticipate that the cost function landscape does not have a funnel-like shape, but may have a golf-course-like shape, in which the configuration space that has a low-cost function is narrow only around the global minimum. Outside this narrow region, the landscape has some noise-driven ruggedness with no global bias towards the global minimum. In such a case, the stochastic search does not work well, in general.

## Acknowledgement

We thank Noriyuki Kodera, and Naoya Terahara and Tohru Minamino for giving the HS-AFM data of actin filament and flagellar FlhA_C_, respectively. This work was partly supported by the Japan Science and Technology Agency (JST) grant (JPMJCR1762) (S.T. and Y.M.) and JSPS KAKENHI Grant Number 19J14515 (T.N.).

---

**Fig S1. Noise distributions in the background regions of experimental AFM images.** We used two AFM movies that contains large background regions, FlhA_C_ monomer (Top) and actin filament (Bottom). After extracting background regions manually, we estimated the stage plane by a simple least squares method and then collected the deviation from the estimated stage height. We took the last 5 frames in the AFM movie. By construction, the mean of the noise is almost equal to zero. The standard deviation of the distribution is 0.284 nm in the case of FlhA_C_ and 0.347 nm in the case of actin filament.

**Fig S2. Twin-experiments for myosin.** (A) One of the reference images in the twin-experiment with the 1 nm / 10 degree probe. (B) The image generated from the predicted structure with the cosine similarity-based cost function. (C) One of the reference images in the twin-experiment with the 3 nm / 20 degree probe. (D) The image generated from the predicted structure with the cosine similarity-based cost function. The color bar shows the height in nm (shared by all of the images here). (E-L) The results of twin-experiment with a reference image generated by the 1 nm / 10 degree probe. The row corresponds to the cost function. The leftmost 4 panels (E, G, I, K) show the lowest scores of images. The next 4 panels (F, H, J, L) show the structure-RMSD of the structures with the lowest score. Red vertical dashed lines show the ground-truth probe shape. (M-T) The results of twin-experiment with a reference image generated by the 3 nm / 20 degree probe, shown in the same way as E-L. The cyan vertical dashed lines show the ground-truth probe shape.

**Fig S3. Twin-experiments with actin filament.** The left 2 columns (ABEFIJ) show the result of the prediction from images with 1nm-wide pixels. The right 2 columns (CDGHKL) show the result with 2nm-wide pixels. The top row (A-D) shows the results with the correlation-based cost function. The middle row (E-H) shows the results with the pixel-RMSD. The bottom row (I-L) shows the results with the penalty function.

**Fig S4**. **The stage position is estimated by a simple least square method.** The left panel shows the experimental image of actin filament. The right panel shows the actual data (gray line) and the estimated position of the stage (red line) along the white dotted line in the left panel.

**Fig S5. Further results of the rigid-body fitting to an HS-AFM experimental image of FlhA_C_ ring** (36). (A) The experimental AFM image used. (B) The background regions that are used in the stage position estimation. White regions are ignored. (C) The corrected AFM image in the reference region used in the rigid-body fitting. (D, G, J) The reference image used in the fitting. The same image as C. (E) The resulting image using the correlation coefficient linearly scaled by a factor of 0.704 to minimize pixel-RMSD. (F) The absolute value of difference between D and E. (H) The resulting image using the correlation coefficient uniformly shifted in −0.89 nm along z axis to minimize pixel-RMSD. (I) The absolute value of the difference between G and H. (K) The resulting image shifted 5.0nm along z axis and scaled linearly by a factor of 0.38 to minimize pixel-RMSD (L) The absolute value of the difference between J and K. (M) The result using the cosine similarity-based cost function. (N) The absolute values of the difference between the reference image (panel C) and the pseudo-AFM image of the best-fit model with the cosine similarity (panel M). (O) The result using the pixel-RMSD. (P) The absolute values of the difference between the reference image (panel C) and the pseudo-AFM image of the best-fit model with the pixel-RMSD (panel O).

**Fig S6. Blind tip reconstruction from pseudo-AFM images.** (A) Noise-free pseudo-AFM image of the 15-mer actin filament used in the twin-experiment in Fig. 4. The image is generated by a 3 nm / 20 degree probe, with the pixel width 1 nm. No noise is added to the image. (B) Cross sections of tip shapes along the X coordinate. The red line denotes the ground truth of the tip shape, i.e., the tip shape used for generating the pseudo-AFM image. The black line is the estimation by the blind tip reconstruction algorithm. (C) Cross sections of tip shapes along the Y coordinate. (D) Pseudo-AFM image with Gaussian noise. Ten different images are generated by adding spatially independent Gaussian noise with the mean 0 nm and the standard deviation 0.3 nm. One of the ten images is shown. (E) Cross sections of tip shapes along the X coordinate. The red line denotes the ground truth of the tip shape. The black lines show the estimated tip shapes by the blind tip reconstruction from the ten images using the threshold parameter of 0 nm. (F) Cross sections of tip shapes along the Y coordinate. (G) and (H) are estimated tip shapes by using the threshold parameter of 0.3 nm.

**Fig S7. Blind tip reconstruction from real AFM data.** (A) The real AFM image of actin filament analyzed by the blind tip reconstruction. This is the same AFM data as those analyzed in Fig. 5. The stage height is corrected by the fitted surface plane. (B) Cross section of the estimated tip shape along the X coordinate. Blind tip reconstruction is applied by using the threshold parameter of 0 nm. (C) Cross section along the Y coordinate. (D) and (E) are estimated tip shapes by using the threshold parameter of 0.3 nm.

## References

1. Deniz AA, Mukhopadhyay S, Lemke EA. Single-molecule biophysics: At the interface of biology, physics and chemistry. J R Soc Interface. 2008;5(18):15–45.

2. Moerner WE, Fromm DP. Methods of single-molecule fluorescence spectroscopy and microscopy. Rev Sci Instrum. 2003;74(8):3597–619.

3. Mazal H, Haran G. Single-molecule FRET methods to study the dynamics of proteins at work. Curr Opin Biomed Eng [Internet]. 2019;12:8–17. Available from: https://doi.org/10.1016/j.cobme.2019.08.007

4. Bacic L, Sabantsev A, Deindl S. Recent advances in single-molecule fluorescence microscopy render structural biology dynamic. Curr Opin Struct Biol [Internet]. 2020;65:61–8. Available from: https://doi.org/10.1016/j.sbi.2020.05.006

5. Saxton MJ, Jacobson K. Single-particle tracking: Applications to membrane dynamics. Annu Rev Biophys Biomol Struct. 1997;26:373–99.

6. Moffitt JR, Chemla YR, Smith SB, Bustamante C. Recent advances in optical tweezers. Annu Rev Biochem. 2008;77:205–28.

7. Alessandrini A, Facci P. AFM: A versatile tool in biophysics. Meas Sci Technol. 2005;16(6).

8. Ando T, Uchihashi T, Kodera N. High-Speed AFM and Applications to Biomolecular Systems. Annu Rev Biophys [Internet]. 2013;42(1):393–414. Available from: http://www.annualreviews.org/doi/10.1146/annurev-biophys-083012-130324

9. Ando T. High-speed atomic force microscopy and its future prospects. Biophys Rev. 2018;10(2):285–92.

10. Uchihashi T, Iino R, Ando T, Noji H. High-speed atomic force microscopy reveals rotary catalysis of rotorless F 1-ATPase. Science (80-) [Internet]. 2011 Aug 5;333(6043):755–8. Available from: http://www.sciencemag.org/cgi/doi/10.1126/science.1205510

11. Kodera N, Yamamoto D, Ishikawa R, Ando T. Video imaging of walking myosin V by high-speed atomic force microscopy. Nature [Internet]. 2010 Nov 10;468(7320):72–6. Available from: http://dx.doi.org/10.1038/nature09450

12. Shibata M, Yamashita H, Uchihashi T, Kandori H, Ando T. High-speed atomic force microscopy shows dynamic molecular processes in photoactivated bacteriorhodopsin. Nat Nanotechnol [Internet]. 2010;5(3):208–12. Available from: http://dx.doi.org/10.1038/nnano.2010.7

13. Miyagi A, Chipot C, Rangl M, Scheuring S. High-speed atomic force microscopy shows that annexin V stabilizes membranes on the second timescale. Nat Nanotechnol [Internet]. 2016;11(9):783–90. Available from: http://dx.doi.org/10.1038/nnano.2016.89

14. Shibata M, Nishimasu H, Kodera N, Hirano S, Ando T, Uchihashi T, et al. Real-space and real-Time dynamics of CRISPR-Cas9 visualized by high-speed atomic force microscopy. Nat Commun [Internet]. 2017;8(1):1–9. Available from: http://dx.doi.org/10.1038/s41467-017-01466-8

15. Kodera N, Noshiro D, Dora SK, Mori T, Habchi J, Blocquel D, et al. Structural and dynamics analysis of intrinsically disordered proteins by high-speed atomic force microscopy. Nat Nanotechnol [Internet]. 2020; Available from: http://dx.doi.org/10.1038/s41565-020-00798-9

16. Scheuring S, Boudier T, Sturgis JN. From high-resolution AFM topographs to atomic models of supramolecular assemblies. J Struct Biol. 2007;159(2 SPEC. ISS.):268–76.

17. Trinh MH, Odorico M, Pique ME, Teulon JM, Roberts VA, Ten Eyck LF, et al. Computational reconstruction of multidomain proteins using atomic force microscopy data. Structure [Internet]. 2012;20(1):113–20. Available from: http://dx.doi.org/10.1016/j.str.2011.10.023

18. Chaves RC, Pellequer JL, Tramontano A. DockAFM: Benchmarking protein structures by docking under AFM topographs. Bioinformatics. 2013;29(24):3230–1.

19. Chaves RC, Teulon J-M, Odorico M, Parot P, Chen SW, Pellequer J-L. Conformational dynamics of individual antibodies using computational docking and AFM. J Mol Recognit. 2013;26(11):596–604.

20. Dasgupta B, Miyashita O, Tama F. Reconstruction of low-resolution molecular structures from simulated atomic force microscopy images. Biochim Biophys Acta - Gen Subj [Internet]. 2020;1864(2):129420. Available from: https://doi.org/10.1016/j.bbagen.2019.129420

21. Markiewicz P, Goh MC. Atomic Force Microscopy Probe Tip Visualization and Improvement of Images Using a Simple Deconvolution Procedure. Langmuir. 1994;10(1):5–7.

22. Oliva AI, Anguiano E, Denisenko N, Aguilar M, Peña JL. Analysis of scanning tunneling microscopy feedback system: Experimental determination of parameters. Rev Sci Instrum [Internet]. 1995 Aug;66(5):3196–203. Available from: http://aip.scitation.org/doi/10.1063/1.1147077

23. Markiewicz P, Goh MC. Simulation of atomic force microscope tip-sample/sample-tip reconstruction. J Vac Sci Technol B Microelectron Nanom Struct. 1995;13(3):1115–8.

24. Tranchida D, Piccarolo S, Deblieck RAC. Some experimental issues of AFM tip blind estimation: The effect of noise and resolution. Meas Sci Technol. 2006;17(10):2630–6.

25. Trinh MH, Odorico M, Bellanger L, Jacquemond M, Parot P, Pellequer JL. Tobacco mosaic virus as an AFM tip calibrator. J Mol Recognit. 2011;24(3):503–10.

26. Lutter L, Serpell CJ, Tuite MF, Serpell LC, Xue WF. Three-dimensional reconstruction of individual helical nano-filament structures from atomic force microscopy topographs. Biomol Concepts. 2020;11(1):102–15.

27. Niina T, Fuchigami S, Takada S. Flexible Fitting of Biomolecular Structures to Atomic Force Microscopy Images via Biased Molecular Simulations. J Chem Theory Comput. 2020;16(2):1349–58.

28. Orzechowski M, Tama F. Flexible fitting of high-resolution x-ray structures into cryoelectron microscopy maps using biased molecular dynamics simulations. Biophys J. 2008;95(12):5692–705.

29. Fuchigami S, Niina T, Takada S. Bayesian Statistical Inference of Experimental Parameters via Biomolecular Simulations: Atomic Force Microscopy. Front Mol Biosci. 2021;8:56.

30. Ando T, Kodera N, Uchihashi T, Miyagi A, Nakakita R, Yamashita H, et al. High-speed atomic force microscopy for capturing dynamic behavior of protein molecules at work. e-Journal Surf Sci Nanotechnol. 2005;3(December):384–92.

31. Amyot R, Flechsig H. BioAFMviewer: An interactive interface for simulated AFM scanning of biomolecular structures and dynamics. PLoS Comput Biol [Internet]. 2020;16(11):1–12. Available from: http://dx.doi.org/10.1371/journal.pcbi.1008444

32. Niina T, Kato S, Koide H. ToruNiina/afmize [Internet]. zenodo. zenodo; 2020. Available from: https://github.com/ToruNiina/afmize

33. Kubo S, Li W, Takada S. Allosteric conformational change cascade in cytoplasmic dynein revealed by structure-based molecular simulations. PLoS Comput Biol. 2017;13(9):1–27.

34. Sack S, Müller J, Marx A, Thormählen M, Mandelkow EM, Brady ST, et al. X-ray structure of motor and neck domains from rat brain kinesin. Biochemistry. 1997;36(51):16155–65.

35. Gurel PS, Kim LY, Ruijgrok P V., Omabegho T, Bryant Z, Alushin GM. Cryo-EM structures reveal specialization at the myosin VI-actin interface and a mechanism of force sensitivity. Elife. 2017;6(Md):1–33.

36. Terahara N, Inoue Y, Kodera N, Morimoto VY, Uchihashi T, Imada K, et al. Insight into structural remodeling of the FlhA ring responsible for bacterial flagellar type III protein export. Sci Adv. 2018;4(4):eaao7054.

37. Uchihashi T, Kodera N, Ando T. Guide to video recording of structure dynamics and dynamic processes of proteins by high-speed atomic force microscopy. Nat Protoc. 2012;7(6): 1193–206.

38. Katchalski-Katzir E, Shariv I, Eisenstein M, Friesem AA, Aflalo C, Vakser IA. Molecular surface recognition: Determination of geometric fit between proteins and their ligands by correlation techniques. Proc Natl Acad Sci U S A. 1992;89(6):2195–9.

39. Villarrubia JS. Morphological estimation of tip geometry for scanned probe microscopy. Surf Sci. 1994;321(3):287–300.

40. Bakucz P, Krüger-Sehm R, Koenders L. Investigation of blind tip estimation. Rev Sci Instrum. 2008;79(7).

41. Williams PM, Shakesheff KM, Davies MC, Jackson DE, Roberts CJ, Tendler SJB. Blind reconstruction of scanning probe image data. J Vac Sci Technol B Microelectron Nanom Struct [Internet]. 1996 Mar 1;14(2):1557. Available from: http://scitation.aip.org/content/avs/journal/jvstb/14/2/10.1116/1.589138

